# Nicotine affects protein complex rearrangement in *Caenorhabditis elegans* cells

**DOI:** 10.1101/052738

**Authors:** Robert Sobkowiak, Andrzej Zielezinski, Wojciech M. Karlowski, Andrzej Lesicki

## Abstract

Nicotine may affect cell function by rearranging protein complexes. We aimed to determine nicotine-induced alterations of protein complexes in *Caenorhabditis elegans* cells, thereby revealing links between nicotine exposure and protein complex modulation. We compared the proteomic alterations induced by low and high nicotine concentrations (0.01 mM and 1 mM) with the control (no nicotine) *in vivo* by using mass spectrometry (MS)-based techniques, specifically the CTAB discontinuous gel electrophoresis coupled with liquid chromatography (LC-MS/MS and spectral counting. As a result, we identified dozens of *C. elegans* proteins that are present exclusively or in higher abundance in either nicotine-treated or untreated worms. Based on these results, we report a network that captures the key protein components of nicotine-induced protein complexes and speculate how the different protein modules relate to their distinct physiological roles. Using functional annotation of detected proteins, we hypothesize that the identified complexes can modulate the energy metabolism and level of oxidative stress. These proteins can also be involved in modulation of gene expression and may be crucial in Alzheimer’s disease. The findings reported in our study reveal intracellular interactions of many proteins with cytoskeleton and may contribute to the understanding of the mechanisms of nicotinic acetylcholine receptor (nAChR) signaling and trafficking in cells.

**Significance of the study:** Most of us are affected by nicotine, not only because of the common use of tobacco products. Nicotine is also included in many popular vegetables of the family Solanaceae, such as tomatoes and peppers. However, these two sources provide the body with radically different doses of nicotine.

Strong biological effects of nicotine rely on binding to the nicotinic receptor, which partially mimics the action of the natural hormone acetylcholine. In our study we used the nematode *Caenorhabditis elegans* as a model organism. General principles of functioning of human cells and *C. elegans* cells are similar. The worm is, however, a compromise between simplicity and complexity, which may facilitate understanding of the more complex systems, like the human body. Our study revealed in the presence of a low nicotine concentration a different composition of polypeptides in the organism than in the presence of a high nicotine concentration. The rearrangements of protein complexes concern proteins involved e.g. in the course of Alzheimer’s disease, which seems interesting in the context of our aging societies. From the perspective of the development of biological research, the ability to identify the components of large protein complexes can contribute to a better understanding of the functioning of cells.

## 1 Introduction

Nicotine, contained in tobacco, is a common drug of addiction and a leading cause of preventable deaths in developed countries[1]. This alkaloid interacts with a broad population of nicotinic acetylcholine receptors (nAChRs), with varying affinities and downstream effectors [2, 3]. Nicotine competes with the natural hormone acetylcholine for binding to the receptors. Mammals have 17 genes encoding for nAChR subunits, in contrast to the nematode *Caenorhabditis elegans*, which has 29 nAChR subunits [4]. Nicotinic receptors appeared in evolution long before the development of the nervous system. They are found in bacteria, plants, and lower vertebrates to regulate basic cell functions, such as proliferation, differentiation, cytoskeletal organization, cell–cell contact, migration, secretion, and absorption [5]. Therefore, cholinergic signaling appeared in the early stages of evolution and seems to be very conservative.

The free-living nematode *C. elegans* is one of the best-studied multicellular model organisms. *C. elegans* has emerged as a powerful experimental system to study the molecular and cellular aspects of human disease *in vivo*. It has been estimated that about 42% of the human disease genes have an ortholog in the genome of *C. elegans*, including those genes associated with Alzheimer’s disease, Parkinson’s disease, spinal muscular atrophy, hereditary nonpolyposis colon cancer, and many others age-related disorders [6].

Owing to its short lifespan, transparent body, and completed genome sequence, it has become an ideal model system in basic biological research. It is also an excellent model organism for the study of drug action [7, 8], including studies of the response to nicotine [9-12].

After no more than 60 min in the presence of nicotine, most of the worms completely change their behavior. Depending on nicotine concentration, they move faster than in control conditions or are paralyzed [10]. The reaction in some conditions is quick and takes less than 10 min. Therefore, it could not be explained by protein synthesis *de novo*, but rather by a disturbance of cholinergic signaling and dysfunction of motor protein complexes. In *C. elegans*, proteomic approaches are part of the essential toolbox for the study of gene function. It is well known that protein function is often dependent on its interactions with other molecules, by formation of specific, macromolecular protein complexes. Thus the same proteins may be present in different types of complexes [13].

Protein interactions can be analyzed by different genetic, biochemical, and physical methods (for reviews, see [14, 15]). Some techniques enable high-throughput screening of a large number of proteins in a cell, such as yeast two-hybrid, tandem affinity purification, mass spectrometry (MS), protein microarrays, synthetic lethality, and phage display [13]. Other methods focus on monitoring and characterizing specific biochemical and physicochemical properties of a protein complex. In the *C. elegans* community, the MS-based proteomic studies of the protein–protein interactions are becoming increasingly common [13].

Four proteomic studies have considerably expanded the number of proteins predicted to be associated with nAChRs [16-19]. Kabbani and colleagues used matrix-assisted laser desorption ionization time-of-flight tandem MS to identify 21 proteins that were either pulled down with the TM3-TM4 loop of mouse β2 or co-immunoprecipitated with β2 [16]. Paulo et al. used MS to analyze brain proteins, affinity-immobilized by α-bungarotoxin (which binds to α7 nAChRs), from wild-type mice and from mutant mice lacking α7 [17]. A comparison of the results identified 55 proteins in wild-type samples that were not present in comparable brain samples from α7 nAChR knockout mice [17]. Paulo et al. compared the proteomic alterations induced by nicotine treatment in cultured pancreatic cells by using MS-based techniques, specifically SDS-PAGE coupled with liquid chromatography (LC)-MS/MS and spectral counting [18]. In their study, out of thousands of proteins in pancreatic cells, hundreds were identified exclusively or in higher abundance in either nicotine-treated or untreated cells. Using 2-dimensional electrophoresis, Yeom et al. presented the proteomic analysis of nicotine addiction-associated proteins in the striatum of rat brains [19].

Due to the limited number of studies on nicotine’s effects on rearrangements of protein complexes, we initiated a study on proteomic effects of nicotine on *C. elegans*. Here we report identification and characterization of protein complexes associated with nicotine treatment. For the first time in *C. elegans*, we used the CAT gel system to observe protein complexes, and identified interacting proteins by MS, during the early response to nicotine. The potential role of nicotine in Alzheimer’s disease, insulin signaling, fatty acid metabolism, oxidative stress, hormesis, and chromatin modulation is discussed. To our knowledge, this study for the first time demonstrates a putative role of nicotine in modulation of protein complex composition.

## 2 Experimental Section

### 2.1 Materials

Tricine, cetyltrimethylammonium bromide (CTAB), arginine, and (-)-nicotine (≥99%) (N3876) were purchased from Sigma-Aldrich, St. Louis, MO. Acrylamide and bis-acrylamide were obtained from Fluka, distributed by Sigma-Aldrich, St. Louis, MO.

### 2.2 Caenorhabditis elegans maintenance

All tests were performed on the wild-type Bristol N2 strain of *C. elegans* obtained from the Caenorhabditis Genetics Center (CGC) at the University of Minnesota (USA). Standard methods were used for the maintenance and manipulation of strains [20]. Nematodes were maintained at 22°C on 5 cm nematode growth medium (NGM) agar plates seeded with *Escherichia coli* (OP50).

### 3. Caenorhabditis elegans synchronization

L1 larvae were prepared by egg synchronization. The hermaphrodites were lysed in 20% bleach (Clorox), 0.5 M NaOH, until fragmented. The resulting eggs were incubated in M9 buffer [20] without food overnight (14–18 h) at 22°C with agitation (150 rpm), to allow larvae to hatch, and reach arrested development due to starvation [20]. These larvae were used to obtain synchronous cultures of *C. elegans*. Large quantities of *C. elegans* were grown in 100 mL of liquid S-medium, by using concentrated *E. coli* OP50 as a food source [10, 20]. A synchronous population of worms was needed in order to eliminate variation in results due to age differences.

### 2.4 Experimental workflow

The 100-mL liquid cultures of *C. elegans* were divided into three equal parts. The first part was collected as a control (no nicotine), the second was challenged on 1 mM (–)-nicotine for 60 min, and the third one was treated with 0.01 mM (–)-nicotine for 60 min. The experimental workflow is summarized in Figure 1. The experiments were performed in parallel and we repeated the experiment (four independent experiments) for both the nicotine-exposed and control groups.

**Figure 1.**
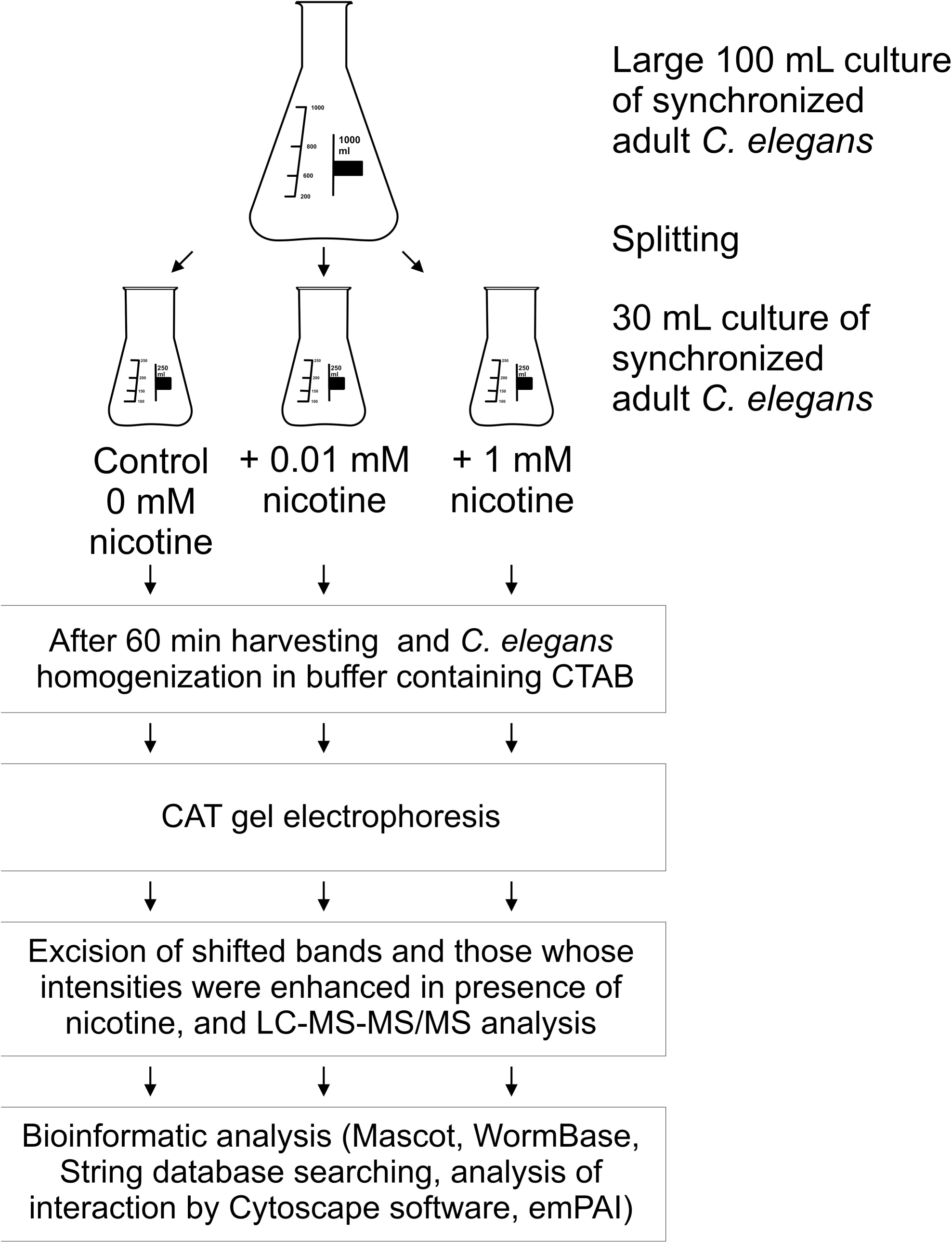
Experimental workflow: adult *Caenorhabditis elegans* were incubated for 60 min with 0.01 mM and 1 mM nicotine or without nicotine (control), next the worms were harvested by centrifugation and protein complexes were fractionated by CAT gel electrophoresis, and finally liquid chromatography coupled to tandem mass spectrometry (LC-MS-MS/MS) analysis was performed, and bioinformatic methods were applied: the Mascot, WormBase, and String database searching, estimation of exponentially modified protein abundance index (emPAI), and interaction analysis with Cytoscape.

### 2.5 Protein extraction

Protein samples were prepared at room temperature immediately prior to loading the gel. The one hundred mg of animals were homogenized with a pestle for about 1 min in 1.5 ml microcentrifuge tubes in 900 μL of CAT sample buffer (10 mM Tricine-NaOH pH 8.8, 1% CTAB, 10% glycerol, 5 mM NaF, and 0.01% phenylmethanesulfonyl fluoride) [21]. The samples were spun in a microfuge for 0.5 min at 16,000 *g* to pellet debris or insoluble material prior to loading the gel. Protein concentrations were determined using the bicinchoninic acid assay (Pierce USA).

### 2.6 CAT gel analysis

The proteins were analyzed by CTAB discontinuous gel electrophoresis (CAT gel system) [21, 22]. A Hoefer SE600 vertical slab gel unit (Hoefer Scientific Instruments, USA) was used with stacking and separating gels consisting of 4 and 10% acrylamide, respectively. The samples containing the same amount of proteins (100 μg of protein per line) were separated at 40 V in CAT gel for 18 h at 22°C.

After the tracking dye had exited the separating gel, the gel was removed and stained with Coomassie blue R-250 (Sigma-Aldrich, St. Louis, MO).

### 2.7 Protein identification

The shifted bands and those whose intensities were markedly enhanced in the presence of nicotine, were excised from the gels. In the control lines, containing the separated extract from *C. elegans* untreated with nicotine, a corresponding area of the gel was excised.

Separation of peptides with high pressure liquid chromatography and subsequent tandem mass spectrometry analysis was performed at the Mass Spectrometry Laboratory of Institute of Biochemistry and Biophysics, Polish Academy of Sciences, Warsaw, Poland. Peptides mixtures were analyzed by LC-MS-MS/MS (liquid chromatography coupled to tandem mass spectrometry) using Nano-Acquity (Waters) LC system and Orbitrap Velos mass spectrometer (Thermo Electron Corp., San Jose, CA). Prior to the analysis, proteins were subjected to standard “in-solution digestion” procedure,[23] during which proteins were reduced with 200 mM dithiothreitol (for 60 minutes at 60°C), alkylated with 200 mM iodoacetamide (45 minutes at room temperature in darkness) and digested overnight with trypsin (sequencing Grade Modified Trypsin - Promega V5111). The peptide mixture was applied to RP-18 precolumn (nanoACQUITY Symmetry^®^ C18 – Waters 186003514) using water containing 0,1% TFA as mobile phase and then transferred to nano-HPLC RP-18 column (nanoACQUITY BEH C18 - Waters 186003545) using an acetonitrile gradient (5 % - 35 % AcN in 70 minutes) in the presence of 0,05% formic acid with the flowrate of 250 nl/min. The column outlet was directly coupled to the ion source of the spectrometer working in the regime of data dependent MS to MS/MS switch. Other MS parameters were the following: m/z range: 300-2000 Th, MS1 resolution: 35 000, MS2 resolution: 15 000, number of precursor ions for fragmentation: 10, intensity threshold for tandem MS: 10^4, precursor selection mass window: 2 Th, charge state screening parameters: 2+ and above, relative CE of 40%, dynamic exclusion: disabled, APEC searching: enabled. A blank run (injection of Milli-Q water) ensuring cross contamination monitoring from previous samples preceded each analysis.

### 2.8 Bioinformatics and data analysis

Acquired raw data were processed by Mascot Distiller (v. 2.4.2) followed by Mascot Search (v. 2.4.1, Matrix Science, London, UK, on-site license) against NCBInr database (v. 20130325, 23 919 380 entries) restricted to Eukaryota. Search parameters for precursor and product ions mass tolerance were 20 ppm and 0.6 Da, respectively, enzyme specificity: trypsin, missed cleavage sites allowed: 1, fixed modification of cysteine by carbamidomethylation and variable modification of lysine carboxymethylation and methionine oxidation.

The peptides with Mascot scores exceeding the threshold value, corresponding to <5% expectation values (in this particular case peptides exceeding Mascot Score of 30, see supplemental data for details) were considered as positively identified. FDR analysis have not been performed, due to relatively small sample size.

All data generated from the gel sections were searched by BLAST against the WormBase database (http://www.wormbase.org/). The interactions among the identified proteins were analyzed using as reference the STRING database of known and predicted protein interactions including both validated physical protein-protein interaction and putative interaction inferred from co-expression, gene fusion, and text mining, etc. (http://string-db.org/) [24]. The open-source software Cytoscape (ver. 3.0.2.) (http://www.cytoscape.org/) was used to visualize molecular interaction networks and biological pathways as well as to integrate these networks with annotations, gene expression profiles and other state data. *C. elegans* proteins were annotated using terms from the Gene Ontology (GO) project (http://www.geneontology.org).

### 2.9 Venn diagrams

The eulerAPE software (drawing area-proportional Euler and Venn diagrams using ellipses) was employed to illustrate the distribution of protein appearance after nicotine treatment, compared to control samples (http://www.eulerdiagrams.org/eulerAPE/).

### 2.10 Estimation of protein abundance

Exponentially modified protein abundance index (emPAI) – an approximate, label-free, relative quantization – was used for estimation of absolute protein abundance in samples [25]. Since emPAI is highly dependent on peptide sampling in MS, only high-quality emPAI values were reported (i.e. containing more than 100 spectra) as described in MASCOT’s emPAI calculation procedure.

## 3 Results

### 3.1 CAT gel electrophoresis of C. elegans proteins in different nicotine concentrations compared to control conditions

We observed explicit differences in the protein band partern on CAT gels (Figure 2). To identify and locate proteins clearly in different lanes, throughout the article the following notation was used: 1 = control worms (no nicotine); 2 = worms exposed to the high concentration of nicotine (1 mM); 3 = worms exposed to the low concentration of nicotine (0.01 mM). Letters indicate the position of bands: a = highest molecular weight; b = intermediate molecular weight; c = lowest molecular weight. As a result, each band is marked by a number followed by letter, e.g. 2c denotes the lowest band in the 1 mM nicotine variant. When comparing the composition of the various bands, e.g. 2a2c means the proteins common to bands 2a and 2c. The number in brackets indicates the number of identified proteins. “Only 2c” means a specific protein, found only in the specified set.

**Figure 2.**
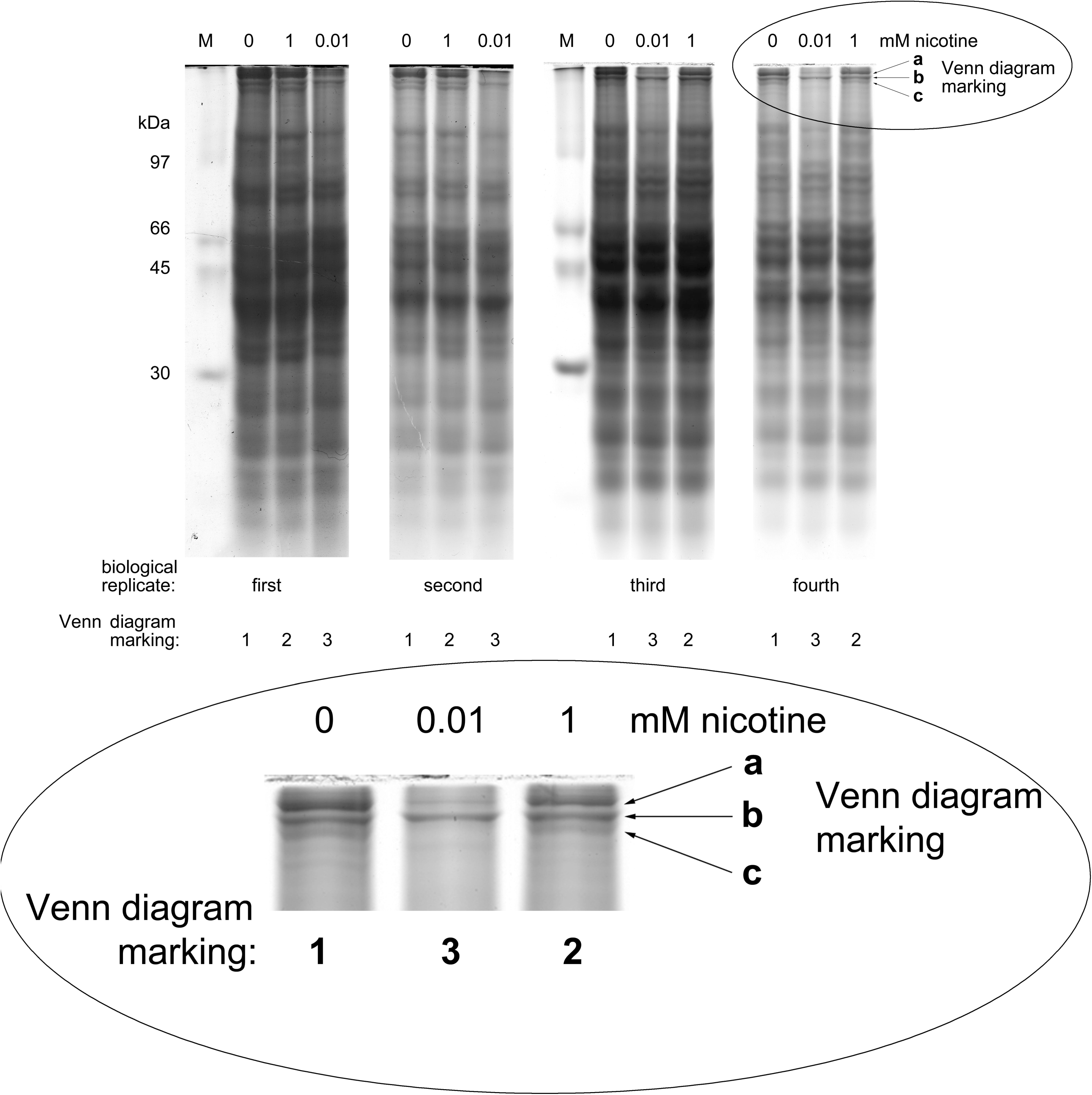
CAT gel electrophoresis of whole adult *Caenorhabditis elegans* lysates. The bands indicated by the arrows were excised and subjected to in-gel digestion followed by mass spectrometric peptide analysis for protein identification. The oval at the bottom shows the enlarged region where reproducible differences were detected and analyzed in worms treated with 0, 0.01, and 1 mM (-)-nicotine. Four biological replicates and nicotine treatment were indicated, with the notation used on Venn diagrams (see Figure 3). 10% CAT gels were applied, electrophoresis duration 18 h, Coomassie blue staining.

**Figure 3.**
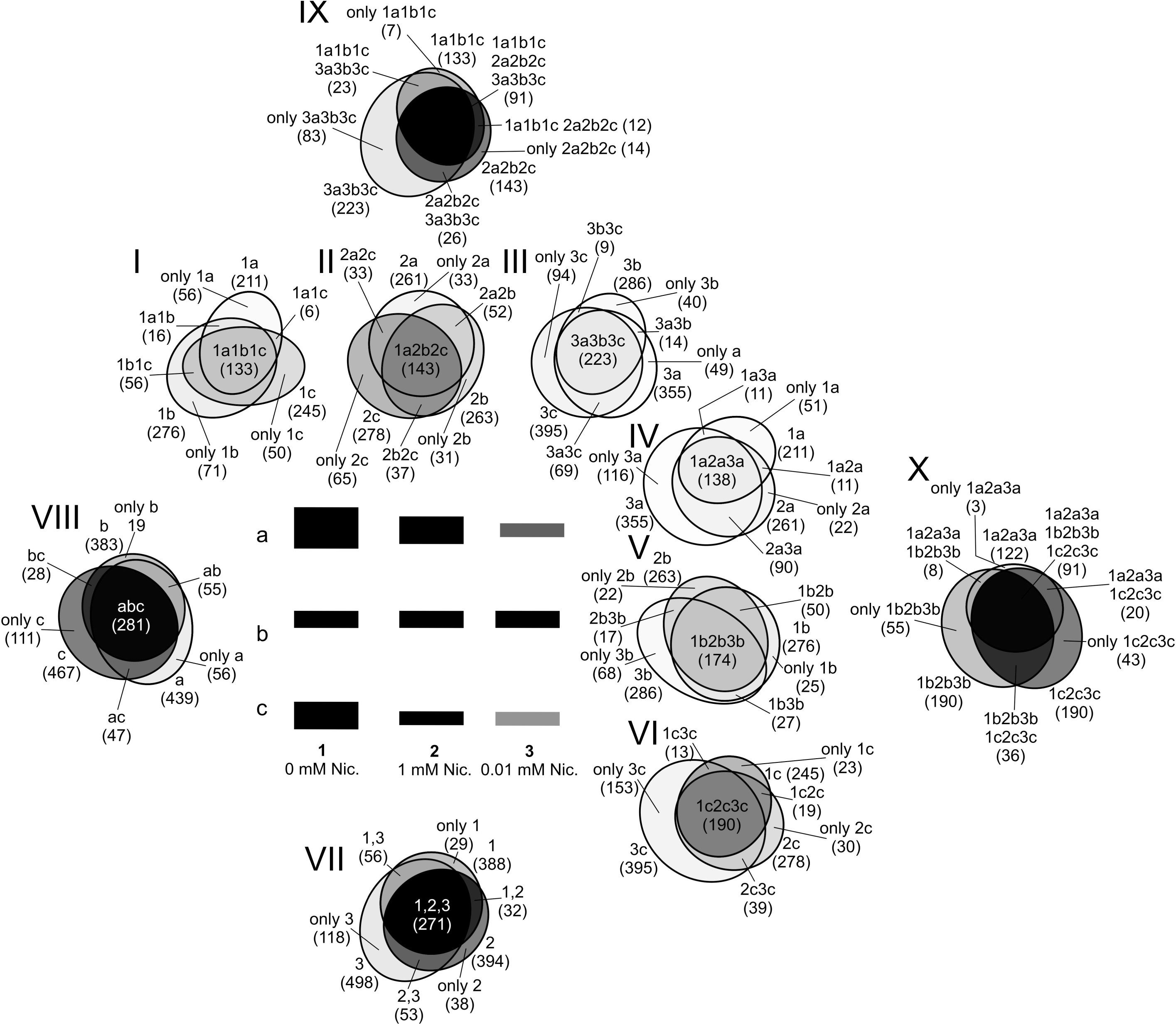
Area-proportional Venn diagrams comparing proteins from *Caenorhabditis elegans* treated with 0.01 mM and 1 mM nicotine and from the control (0 mM nicotine). For example, lalblc means proteins common to the 0 mM nicotine variant, occurring simultaneously in all its bands, while “only la” means proteins unique to the band la only occurring there. The number of proteins detected in each of the sets is given in parentheses. Venn diagrams marked by roman numerals.

Four biological replications were performed. In the gel area where the proteins with the highest mass were located, we observed changes in the position and intensity of the bands in the presence of nicotine, compared to control conditions (see enlarged view in Figure 2). The pattern of bands in the control was more similar to that of the bands from samples treated with 1 mM nicotine than to that of the bands observed in the 0.01 mM variant. The patterns were similar in all the replications of experiments (Figure 2).

### 3.2 Core proteins identified in complexes across all bands in both nicotine-treated and untreated C. elegans

MS analysis identified 597 proteins. We found that approximately 15% *(n =* 91) of the proteins appeared in all gel bands (i.e. 1a, 1b, 1c, 2a, 2b, 2c, 3a, 3b, 3c, see Figures 2 and 3). These proteins, shared among control and both nicotine treatments, are listed in Table 1 (see supporting information). The complexes migrate on CAT gels as three closely spaced bands. Overall, the identified proteins include cytoskeletal components (like actin, myosin, and tubuline), proteins involved in vesicular transport (clathrin, kinesin), and others: spectrins, heat shock proteins, vitellogenin, ribosomal proteins, translation elongation factors, transitional endoplasmic reticulum ATPase (CDC-48), mitochondrial proteins, etc. (Table 1 in supporting information).

### 3.3 Proteins specific to nicotine-treated worms

The proteins unique to the nicotine-treated *C. elegans* cells were compared to the proteins unique to the untreated samples. Generally about 5% of the total proteins identified were exclusive to untreated worms (see Figure 3, Venn VII “only 1”: 29 proteins), while about 6% (see Venn VII “only 2”: 38 proteins) and 20% (see Venn VII “only 3”: 118 proteins) of the total proteins identified were exclusive to high and low nicotine variants, respectively. About 45% (see Venn VII common to 1, 2, and 3: 271 proteins) of the identified proteins were found in both nicotine-treated and untreated worms (Figure 3).

Proteins identified exclusively in nicotine-treated worms are listed in Tables 2, 3, and 4 (see supporting information). We identified 53 proteins unique to both 0.01 mM and 1 mM nicotine-treated worms (Table 2) as well as 118 and 38 proteins unique only to the 0.01 mM nicotine-treated worms (Table 3) and only to the 1 mM nicotine-treated worms (Table 4), respectively. Proteins identified exclusively in untreated worms (control) are listed in Table 5. Proteins unique to “a”, “b”, and “c” bands are shown in Tables 6, 7, and 8, respectively (see supporting information).

### 3.4 Protein complexes

By using Cytoscape and protein interaction data retrieved from the String database, we visualized the interaction networks of the identified proteins in different experimental variants. Most forms of proteins are included in the complex (Figure 4). In Figure 5 we show the core complex of proteins shared by all experimental variants.

**Figure 4.**
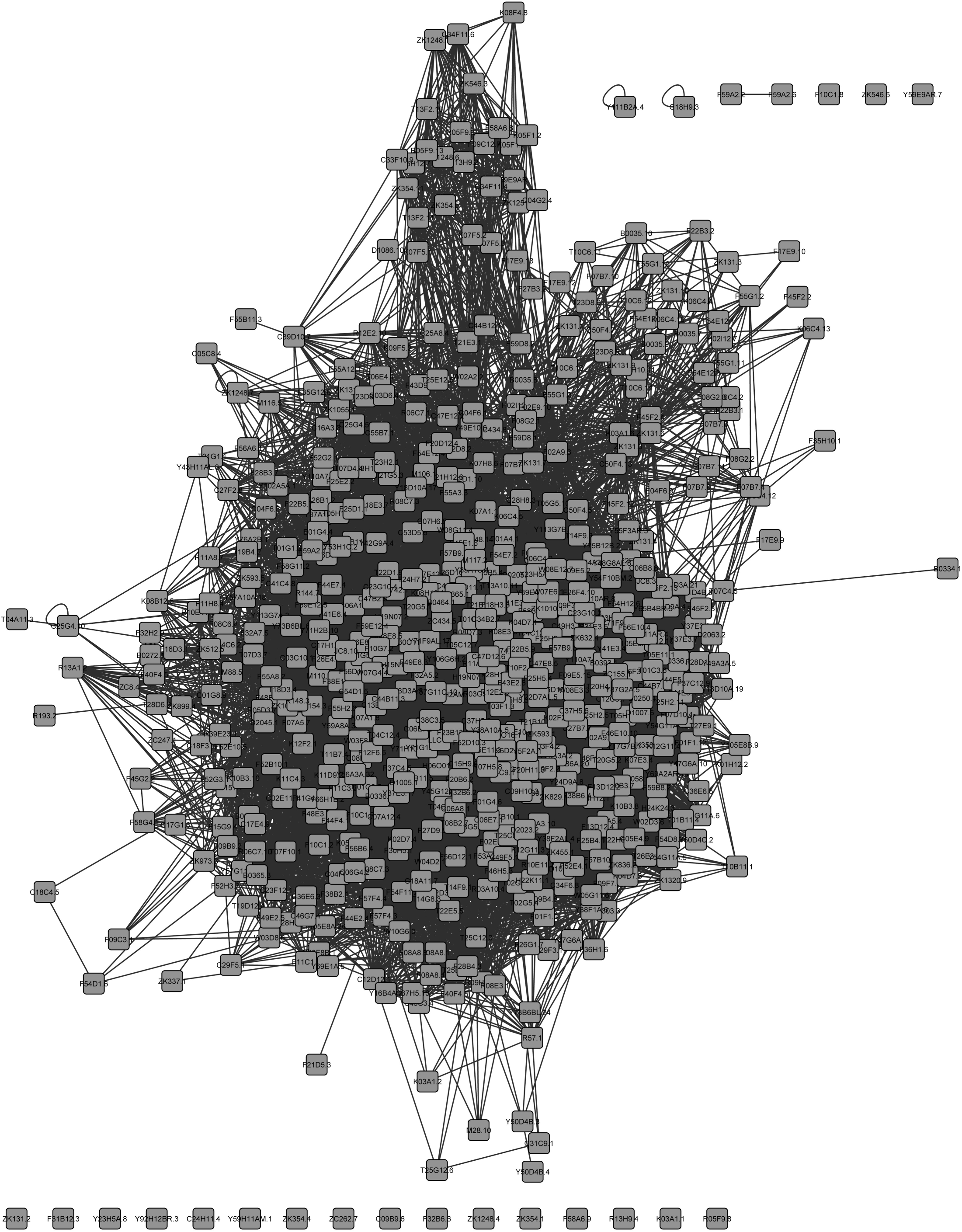
The protein complex network of *Caenorhabditis elegans*: view of the entire identified 597 protein dataset. The black lines represent known protein-protein interactions derived from the String database. For details, see supplemental data.

**Figure 5.**
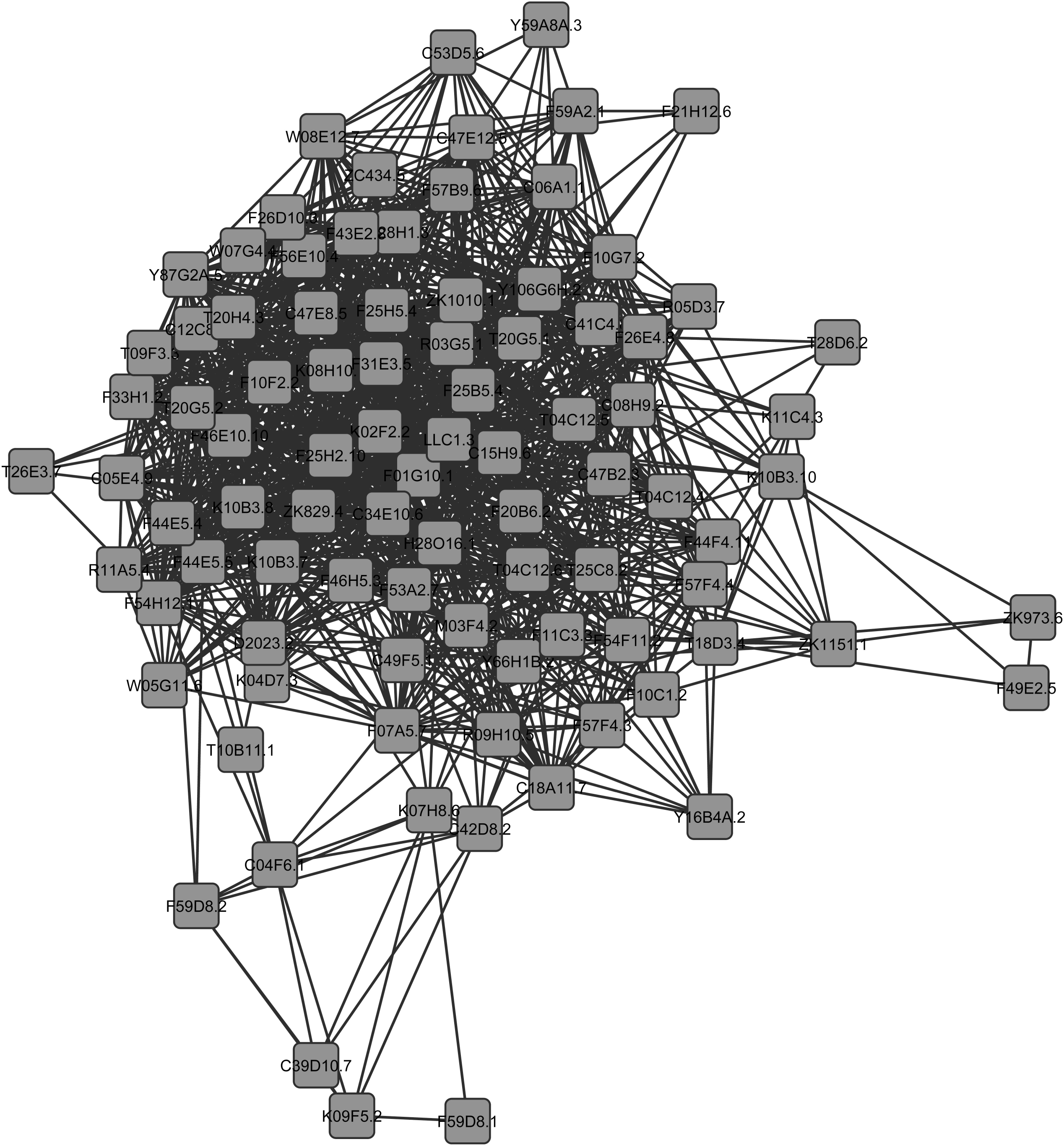
The protein complex network of *Caenorhabditis elegans*: view of the identified core proteins in each band dataset. The black lines represent known protein–protein interactions derived from the String database. The proteins are listed in Table 1. For details, see supplemental data.

### 3.5 Quantitative proteomic differences among all experimental variants

The majority of the proteins identified are common to both nicotine-treated and untreated *C. elegans* cells. Quantitative proteomic analysis was performed to investigate differences in the abundance of these proteins. The protein patterns differed among the control (0 mM nicotine) and two nicotine concentrations (0.01, and 1 mM) (Figure 3). The differences in staining intensity (Figure 2) reflect the relative abundance of proteins. MS analysis revealed dozens of proteins that were present in significantly different relative abundances in the nicotine-treated worms than in the control (see supporting information). In Table 9, we list the differentially abundant proteins that were identified in all experimental variants. While dimethylglycine dehydrogenase was more abundant in the nicotine-treated worms, the remaining proteins of this group were more abundant in the control. Many of the listed proteins – such as profilin, filamin, myosin, and plectin – are common cytoskeletal proteins that bind actin, which may be associated with the protein pattern reorganization observed after nicotine treatment, as shown in Figure 2.

## 4 Discussion

### 4.1 General remarks

We investigated nicotine-induced changes in protein complex patterns in cells of *Caenorhabditis elegans*. Although several proteome studies of *C. elegans*, using various methods of separation, detection, and quantification were reported earlier (for a review, see [26]), we decided to use a different, yet powerful and flexible technique for the separation of proteins. To our best knowledge the CAT electrophoresis and mass spectrometry (MS) was used for the first time to identify protein complexes. The protocol is based on the use of the cationic detergent CTAB [27], which is the most effective agent for the solubilization of highly hydrophobic proteins and integral membrane proteins [28]. CAT electrophoresis allows the fine separation of proteins to be carried out with the retention of native activity [27]. Many proteins separated on CAT gels may remain in native complexes, and under the conditions presented in our study, CTAB may be considered a nondenaturing detergent. The combination of CAT gel electrophoresis and tandem MS is a simple method for revealing many potential interacting proteins associated directly or indirectly with a protein of interest. The use of CAT electrophoresis allowed comparison of the number and intensity of bands in the control versus nicotine-treated nematodes.

Since the isolation of protein complexes from native tissue enables the analysis of interactions within the normal cellular milieu, it provides a significant advantage over traditional protein interaction screens, such as the yeast two-hybrid and tandem affinity purification. The interaction discovery techniques, like the two-hybrid system or affinity purifications, suggest physical associations between proteins [29]. On the other hand, direct physical interactions of proteins might be rather limited, covering perhaps <1% of the theoretically possible interaction space [24]. Proteins do not necessarily need to undergo a stable direct interaction to have a specific, functional interplay. Together with direct, physical interactions, such indirect interactions constitute the larger superset of functional protein–protein associations [24].

We observed major visible differences in the protein-banding patterns on gels, which are not clearly noticeable in the presence of nicotine in other proteomic analyses [18]. Our MS analysis of proteins separated by CAT gel electrophoresis has identified many proteins that were not among the proteins listed by other authors [16-18]. Among the identified peptides, there are also many novel proteins, often referred to as “unnamed”, that have not yet been described in the context of cellular response to nicotine.

Under physiological conditions, the cell is exposed to a number of external factors, of which only a few induce adaptive responses. The process begins at the moment in which the cell membrane protein receptor receives an extracellular signal and begins transduction of the signal across the plasma membrane, and the final cell reaction is initiated. An early response of cells to a signal is mainly based on a change in the activity of the enzymes and protein complex rearrangements. It is known from yeast studies that most of these complexes are composed of several core proteins and several proteins that may temporarily connect to them [30].

The bioinformatic analysis, using existing databases, showed that many of the proteins identified by us form complex networks with permanent interactions (see Figure 5). We suppose that the retention of native protein activity noted by Akins et al. is at least partially due to remaining of proteins in native complexes [27]. The existing data contained in large databases of protein interactions, such as String [24], confirm this hypothesis. Almost all proteins identified by us in the bands interact with each other, suggesting that indeed they may form complexes (see Figures 4 and 5). The ability to detect native protein bindings or associations greatly enhances the ability of researchers to analyze protein complexes. In our experiments we identified the core proteins, present in all tested bands. The core complex encompasses 91 proteins (Table 1). The complex binds other proteins and is involved in many processes. In our opinion, it is most important to discuss here the aspects of cell stress and changes in proteins involved in Alzheimer’s disease.

### 4.2 Nicotine and cell stress

It is well known that nicotine may influence the fat stores, as an increase in body weight is observed in many people after they quit smoking. Besides, it is well established that smoking stimulates lipolysis *in vivo* [31]. Lipolysis appears to be the most sensitive system to nicotine stimulation among various metabolic processes [31]. Nicotine can also stimulate lipolysis directly, by binding to nicotinic cholinergic receptors in adipose tissue [31]. The increase in the rate of lipolysis leads to increased concentrations of free fatty acids. In humans, nicotine decreases food intake and increases thermogenesis, increases the delivery of free fatty acids to the liver and skeletal muscle, and decreases muscle glucose uptake. These effects of nicotine are associated with increased lipoprotein secretion and intramyocellular lipid saturation as well as peripheral insulin resistance [32]. In the presence of 0.01 mM nicotine we found a 4-5-fold decrease in vitellogenin (vit-1 and vit-5), which is predicted to function as a lipid transport protein [33].

Among the proteins unique to the samples treated with the low concentration of nicotine, we found protein phosphatase PPM-1. According to WormBase, the best human ortholog for this protein is protein phosphatase 1 B (PTP1B). PTP1B has emerged as a key regulator of signaling networks that are implicated in metabolic regulation, in particular, insulin signaling, endoplasmic reticulum stress response, cell–cell communication, energy balance, and vesicle trafficking. Thus protein-tyrosine phosphatase 1B is involved in metabolic diseases, such as obesity and type 2 diabetes [34].

In the presence of the low nicotine concentration we found additionally protein phosphatase 2A (PPTR-1). This phosphatase also connects nutrient levels to stress, metabolism, development, longevity, and behavior, specifically through the insulin/IGF-1 signaling pathway [35]. A PPTR-1 regulatory subunit regulates *C. elegans* insulin/IGF-1 signaling by modulating AKT-1 phosphorylation [36]. AKT-1 is a crucial element in signaling of nAChRs [2, 3]. Changes in PPTR-1 levels affect the activity of AKT-1. Activation of Akt by nicotine occurred within minutes at concentrations achievable by smokers [37]. PPTR-1 also regulates DAF-16-dependent outputs of the insulin/IGF-1 signaling pathway, such as fat storage [36]. We observed fatty acid-binding proteins (FAR-6, FAR-1), very low-density lipoprotein receptor (F44E2.4), and many mitochondrial proteins. Nicotine probably contributes to an increase in the demand for ATP produced by mitochondria, which are transported by kinesin to the areas with an increased demand for energy. ATP can be consumed by the ABC transporter SPG-7 metalloprotease, ATP-dependent RNA helicase, and vacuolar H ATPase, which we found in the presence of 0.01 mM nicotine.

In the samples treated with 0.01 mM nicotine we found a 7-fold decrease in talin-1 (TLN-1), as compared to the control conditions. TLN-1 is predicted to have insulin receptor-binding activity, actin-binding activity, and is predicted to be a structural constituent of cytoskeleton.

Among the many proteins we found also acyl-coenzyme A oxidase and 3-ketoacyl-CoA thiolase, which are involved in degradation pathways, such as fatty acid beta-oxidation. Two substrates of acyl-CoA oxidase are acyl-CoA and O_2_, whereas its two products are trans-2,3-dehydroacyl-CoA and H_2_O_2_. Reactive oxygen species (ROS), e.g. H_2_O_2_, are assumed to be detrimental to many biological processes. Catalase (CTL) is a very important enzyme in protecting the cell from oxidative damage by ROS, decomposing H_2_O_2_. In our experiments CTL joined within no more than 60 min to the complex core, and was present in the complex only after nicotine treatment. This suggests a very fast and efficient cell response to the presence of nicotine. CTL activity was found to be significantly increased in nematodes, with restriction of glucose metabolism only after several days [38]. In the presence of nicotine we found all of three *C. elegans* CTLs (CTL-1, CTL-2, and CTL-3).

The increase in fatty acid beta-oxidation is followed by an increase in CTL binding/activity, and results in leveling up oxidative stress resistance [38]. Induction of mitochondrial metabolism might cause a positive response to increased formation of ROS and other stressors, leading to a secondary increase in stress defense, cumulating in reduced net stress levels [38]. Schulz et al.’s (2007) findings are in accord with the hormesis theory of aging, hormesis being defined as a short-lasting and nonlethal stressor that induces the stress response mechanisms of an organism and thereby increases not only stress resistance [38]. Also, hormesis has been observed in *C. elegans* under different conditions of stress, such as direct oxidative and thermal stress [39]. This is in line with our previous work: we found that nicotine in some concentrations is genoprotective and in others it is neutral or genotoxic [11, 40]. We observed in human leukocytes and *C. elegans* cells that treatment with a low concentration of nicotine results in a lower level of DNA damage than in control conditions. We therefore hypothesize that it was due to the presence of CTL in core proteins in places where hydrogen peroxide was generated. High concentrations of nicotine limited the capabilities of cells to defend against ROS and induced damage greater than under control conditions. The low concentrations of nicotine efficiently stimulated the cells’ ROS defense mechanism, as the amount of ROS was even smaller than in control conditions, resulting in a reduced amount of DNA damage in the presence of the low nicotine concentration, as compared to the control [11, 40].

Another symptom of stress was the appearance of proteins that are produced by cells in response to exposure to stressful conditions. Interestingly, the heat shock protein hsp-60 (HSP60) was about 7.5-fold more abundant at the low nicotine concentration in comparison to the control or high nicotine concentration. HSP60 is a mitochondrial chaperonin that is typically held responsible for the transportation and refolding of proteins from the cytoplasm into the mitochondrial matrix. It has also been shown to be involved in stress response. The heat shock response is a homeostatic mechanism that protects cells from damage. Calabrase et. al.’s data are in favor of the importance of HSP-60 in the heat shock signal pathway as a basic mechanism of defense against neurotoxicity elicited by free radical oxygen produced in neurodegenerative disorders [41].

### 4.3 Nicotine and proteins involved in Alzheimer’s disease

The most visible differences in the protein complex partern were observed in the presence of low nicotine concentrations, compared to the controlled conditions and high nicotine concentration (Figure 2). In the presence of 0.01 mM nicotine, a lower intensity of bands was observed. This may be due to the fact that in the analyzed protein complexes several peptidases were found, e.g. NEPrilysin metallopeptidase, aspartyl protease (ASP-1, ASP-2), and elements of proteasome RPN-10. Surprisingly, a plethora of other proteins unique to the low nicotine concentration associated with the core complex described above was revealed.

Nicotine treatment, by stimulation of the brains of Alzheimer’s disease patients, has been shown to attenuate the decline in some of the cognitive deficits symptomatic of the disease and particularly effective in reversing attentional deficits [42]. Amyloid beta (Aβ) is a peptide whose misfolding and aggregation in abnormal neural tissue has been implicated as a cause of Alzheimer’s disease. The following proteinases have the ability to degrade the Aβ peptide in Alzheimer’s disease: NEPrilysin, insulin-degrading enzyme, plasmin, endothelin converting enzymes, cathepsin, glutamate carboxypeptidase II, angiotensin-converting enzyme, etc. [43, 44]. Deficiency of NEPrilysin and insulin-degrading enzyme caused increased cerebral accumulation of endogenous Aβ in transgenic models of Alzheimer’s disease *in vivo* [44]. In our experiment in the presence of a low concentration of nicotine we found most probably the same proteins from the NEPrilysin metallopeptidase family: nep-22, nep-2, F44E7.4, and cathepsin. These findings seem to confirm the beneficial effects of nicotine in the course of Alzheimer’s disease treatment. They may be additionally supported by the fact that we found also F44E2.4. According to WormBase its best human ortholog is a very low-density lipoprotein receptor. Interestingly, low-density lipoprotein receptor-related protein (LRP) regulates Aβ clearance [44]. In the pathogenesis of Alzheimer’s disease a critical role may be played also by prion protein [45]. The cellular form of the prion protein interacts with and inhibits β-secretase (BACE1), the rate-limiting enzyme in the production of Aβ [45]. β-secretase is a kind of aspartyl protease. In the presence of 0.01 mM we found prion-like proteins PQN-85, PQN-27, and aspartyl protease (ASP-1 and ASP-3). In all experimental variants (0, 0.01, and 1 mM nicotine) we observed prion-like-(Q/N-rich)-domain-bearing protein (PQN-87). However in the presence of 0.01 mM nicotine a decrease was observed in the abundance of this protein to 64%, as compared to control conditions. This strengthens the relationships of nicotine with prevention of Alzheimer’s disease. In the presence of both concentrations of nicotine, we found transthyretin-like protein (TTR-1). Transthyretin is thought to have beneficial side effects, by binding to the beta-amyloid protein, thereby preventing beta-amyloid’s natural tendency to accumulate into the plaques associated with the early stages of Alzheimer’s disease [44]. Failures in the treatment of Alzheimer’s disease by using nicotine may result from the fact that in the presence of 1 mM nicotine we observed only a 27% increase in the abundance of this protein, as compared to the control. Additionally, in the presence of 0.01 mM nicotine we found TTR-18 and TTR-16, which are transthyretin-like proteins. Transthyretin (TTR) is the carrier of the thyroid hormone thyroxine. A single *in vivo* study in *Caenorhabditis elegans* suggested that wild-type human TTR could suppress the abnormalities seen when Aβ was expressed in the muscle cells of the worm [46].

Our results may indicate that indeed nicotine contributes to the emergence of Alzheimer’s-related peptidase and prion-like proteins in the complex, and that these proteins are involved in the response to alkaloid. Thus nicotinic drug treatment probably should be taken into account as a putative novel protective therapy in Alzheimer’s disease, but in a strictly controlled dose, suited to a given patient.

### 4.4 Other effects of nicotine on protein complexes

Binding of nicotine to the nicotine receptor starts the signal transduction process. It has been shown, using confocal fluorescence microscopy and immunogold labeling in electron microscopic studies, that actin is co-localized with the nAChR, which is regarded as a crucial element in the response to nicotine [47]. Results from *in vitro* cotransfection experiments suggest that acetylcholine receptors are tethered to the actin cytoskeleton via several adapter proteins [48]. Here it is worth noting that among the proteins unique to the “c” band (see supporting information Table 8), we found in all experimental variants (0, 0.01, 1 mM nicotine concentrations) the NRA-4 nicotinic receptor-associated protein. NRA-4 may control nAChR subunit composition or allow only certain receptor assemblies to leave the endoplasmic reticulum [49]. Only in the presence of low nicotine concentrations we found acetylcholine receptor ACR-21, which may indicate that only in such circumstances the nicotinic receptor is physically connected to the protein complex observed by us. We hypothesize that the nicotinic receptor is not always directly connected to the protein complex involved in the response to nicotine. This may explain some different results of experiments designed to detect proteins that form a complex with the nicotinic receptor [16, 17].

In our experiment we found also seventeen of histone H3 forms (see supporting information Table 4). Nucleosome remodeling is associated with histone modifications. It is noteworthy that we have identified 17 different histone H3 variants, showing highest similarity to human histone H3.2, deposited in transcriptionally silent areas that can be reversibly activated [50]. Besides, histone H3 is the most extensively modified of the five histones. It is an important protein in the field of epigenetics, where its sequence variants and variable modification states are thought to play a role in the dynamics and long-term regulation of genes. Alterations in gene expression are implicated in the pathogenesis of several neuropsychiatric disorders, including drug addiction. Increasing evidence indicates that changes in gene expression in neurons, in the context of animal models of addiction, are mediated in part by epigenetic mechanisms that alter chromatin structure on specific gene promoters [51]. Exposure to drugs of abuse induces changes within the brain’s reward regions in several modes of epigenetic regulation – histone modifications such as acetylation and methylation [52]. A recent genetic screening strategy has also identified histone H3 lysine 9 (H3K9) methylation as a key determinant for peripheral tethering of silenced heterochromatin at the nuclear envelope [53]. Histones were observed by us in the protein complex in the presence of 1 mM nicotine. In the presence of all nicotine concentrations we discovered histone-arginine N-methyltransferase (PRMT-1) and S-adenosylmethionine (SAM). SAM generates the universal donor for methylation reactions in eukaryotic cells. We also found in both concentrations of nicotine the protein arginine N-methyltransferase that functions as a histone methyltransferase specific for histone H4. Interestingly, we found histone H4 in the 0.01 mM nicotine variant at the same level as in the control and two-fold lower than in the presence of 1 mM of nicotine.

The nucleosome remodeling described above may explain our previous observation that nicotine induces an increase in the size of the comet head that is formed in the comet assay in place of the nucleus [11, 40]. The increase in head radius was probably due to forming a DNA bulge. This observation was made in the low concentration of nicotine both in *C. elegans* cells and in human leucocytes [11, 40].

Also it should be stressed that our results indicate an about 23-fold and 7.7-fold increase in putative mitochondrial dimethylglycine dehydrogenase in the presence of 0.01 mM and 1 mM nicotine, respectively. Dimethylglycine dehydrogenase is a mitochondrial matrix enzyme involved in the metabolism of choline, converting dimethylglycine to sarcosine [54]. We suppose that nicotine induces imbalance in cholinergic signalization by stimulation of only the nAChR branch. Activation of only half of the cholinergic pathway by nicotine is detected by cells and in response the cells remove choline – the substrate used for the synthesis of acetylcholine. This is a probable explanation of the huge increase in the quantity of dimethylglycine dehydrogenase.

### 4.5 Conclusions

A good understanding of the remodeling of protein complexes will make it possible to create a map showing the interactions between all or some of the proteins of the proteome. This will be an important step in combining proteomics with cell biology. This process resembles constructing a 3-dimensional puzzle composed of several 2-dimensional puzzles stacked on one another, requiring the assembly of, often oddly shaped, interlocking pieces. This may be of great importance for the understanding of the functioning of a single cell under normal conditions, and under conditions of various diseases or drug dependence. Thus, the associated protein complexes, in addition to nAChRs, might represent targets for the development of drugs to treat various diseases, like Alzheimer’s disease, obesity or diabetes. There is increasing interest in finding small molecules, e.g. nicotine and its derivatives, that affect protein–protein interactions, and this strategy could be used to identify the compounds that either disrupt or strengthen interactions between nAChRs and an associated protein as part of developing novel therapeutics. We have discovered a number of unnamed proteins in the complex core and in the presence of the low nicotine concentration, of which almost nothing is known. The role of these proteins in the cell’s response to nicotine should be taken into account and needs to be confirmed in future experiments. Verification of the proteins that we identified as a potential nicotine response interactome will need to be performed via traditional biochemical methods, such as Western blotting and coimmunoprecipitation, before more concerted efforts are devoted to the detailed study of these interactions. The detected proteins may be useful in revealing the molecular mechanism of nicotine addiction and in isolating the potential drug targets for nicotine addiction-associated disease therapy. The compiled list of proteins in the interaction network obtained from this study is not meant to be comprehensive, nor have we validated the identified proteins via orthogonal assays. Nevertheless, we think that our work has laid a solid foundation for further exploration of the nicotine response interactome in both neuronal and non-neuronal tissues. Furthermore, the CAT electrophoresis, together with the proteomic MS analysis procedure, could also be applied to the investigation of the multiprotein complexes.

## Acknowledgements

Authors thank Dr. Janusz Dębski (Mass Spectrometry Laboratory, Institute of Biochemistry and Biophysics, Warsaw, Poland) for his assistance with mass spectrometry analysis. We thank Dr. R Rucinska-Sobkowiak for critical reading of this manuscript. We thank also Sylwia Ufhalska, an author’s editor, for help in improving the manuscript.

## References

[1] WHO, WHO global report: mortality attributable to tobacco. http://www.who.int/tobacco/publications/surveillance/rep_mortality_attributable/en/ 2012.

[2] Schuller, H. M., Is cancer triggered by altered signaling of nicotinic acetylcholine receptors? Nat Rev Cancer 2009, 9, 195–205.

[3] Sobkowiak, R., Lesicki, A., Cell signaling pathways activated by nicotine. Advances in Cell Biology 2011, 38, 581–596.

[4] Jones, A. K., Davis, P., Hodgkin, J., Sattelle, D. B., The nicotinic acetylcholine receptor gene family of the nematode *Caenorhabditis elegans*: an update on nomenclature. Invert Neurosci 2007, 7, 129–131.

[5] Skok, M. V., Editorial: To channel or not to channel? Functioning of nicotinic acetylcholine receptors in leukocytes. J Leukoc Biol 2009, 86, 1–3.

[6] Markaki, M., Tavernarakis, N., Modeling human diseases in *Caenorhabditis elegans*. Biotechnol J 2010, 5, 1261–1276.

[7] Schafer, W. R., Addiction research in a simple animal model: the nematode Caenorhabditis elegans. Neuropharmacology 2004, 47, 123–131.

[8] Wolf, F. W., Heberlein, U., Invertebrate models of drug abuse. J Neurobiol 2003, 54, 161–178.

[9] Feng, Z., Li, W., Ward, A., Piggott, B. J., et al., A *C. elegans* model of nicotine-dependent behavior: regulation by TRP-family channels. Cell 2006, 127, 621–633.

[10] Sobkowiak, R., Kowalski, M., Lesicki, A., Concentration- and time-dependent behavioral changes in *Caenorhabditis elegans* after exposure to nicotine. Pharmacol Biochem Behav 2011, 99, 365–370.

[11] Sobkowiak, R., Lesicki, A., Genotoxicity of nicotine in cell culture of *Caenorhabditis elegans* evaluated by the comet assay. Drug Chem Toxicol 2009, 32, 252–257.

[12] Wonnacott, S., Sidhpura, N., Balfour, D. J., Nicotine: from molecular mechanisms to behaviour. Curr Opin Pharmacol 2005, 5, 53–59.

[13] Fonslow, B. R., Moresco, J. J., Tu, P. G., Aalto, A. P., et al., Mass spectrometry-based shotgun proteomic analysis of *C. elegans* protein complexes. WormBook 2014, 1–18.

[14] Shoemaker, B. A., Panchenko, A. R., Deciphering protein-protein interactions. Part I. Experimental techniques and databases. PLoS Comput Biol 2007, 3, e42.

[15] Shoemaker, B. A., Panchenko, A. R., Deciphering protein-protein interactions. Part II. Computational methods to predict protein and domain interaction partners. PLoS Comput Biol 2007, 3, e43.

[16] Kabbani, N., Woll, M. P., Levenson, R., Lindstrom, J. M., Changeux, J. P., Intracellular complexes of the beta2 subunit of the nicotinic acetylcholine receptor in brain identified by proteomics. Proc Natl Acad Sci U S A 2007, 104, 20570–20575.

[17] Paulo, J. A., Brucker, W. J., Hawrot, E., Proteomic analysis of an alpha7 nicotinic acetylcholine receptor interactome. J Proteome Res 2009, 8, 1849–1858.

[18] Paulo, J. A., Urrutia, R., Kadiyala, V., Banks, P., et al., Cross-species analysis of nicotine-induced proteomic alterations in pancreatic cells. Proteomics 2013, 13, 1499–1512.

[19] Yeom, M., Shim, I., Lee, H.-J., Hahm, D.-H., Proteomic analysis of nicotine-associated protein expression in the striatum of repeated nicotine-treated rats. Biochemical and Biophysical Research Communications 2005, 326, 321–328.

[20] Stiernagle, T., Maintenance of C. elegans. WormBook 2006, 1–11.

[21] Walker, J. M., The Protein Protocols Handbook, Humana Press Inc., Totowa, New Jersey 2002.

[22] Akins, R. E., Tuan, R. S., Separation of proteins using cetyltrimethylammonium bromide discontinuous gel electrophoresis. Mol Biotechnol 1994, 1, 211–228.

[23] Przewloka, M. R., Venkei, Z., Bolanos-Garcia, V. M., Debski, J., et al., CENP-C is a structural platform for kinetochore assembly. Curr Biol 2011, 21, 399–405.

[24] Franceschini, A., Szklarczyk, D., Frankild, S., Kuhn, M., et al., STRING v9.1: protein-protein interaction networks, with increased coverage and integration. Nucleic Acids Res 2013, 41, D808–815.

[25] Ishihama, Y., Oda, Y., Tabata, T., Sato, T., et al., Exponentially modified protein abundance index (emPAI) for estimation of absolute protein amount in proteomics by the number of sequenced peptides per protein. Mol Cell Proteomics 2005, 4, 1265–1272.

[26] Schrimpf, S. P., Hengartner, M. O., A worm rich in protein: Quantitative, differential, and global proteomics in *Caenorhabditis elegans*. J Proteomics 2010, 73, 2186–2197.

[27] Akins, R. E., Levin, P. M., Tuan, R. S., Cetyltrimethylammonium bromide discontinuous gel electrophoresis: Mr-based separation of proteins with retention of enzymatic activity. Anal Biochem 1992, 202, 172–178.

[28] Polati, R., Zapparoli, G., Giudici, P., Bossi, A., A CTAB based method for the preparation of total protein extract of wine spoilage microrganisms for proteomic analysis. J Chromatogr B Analyt Technol Biomed Life Sci 2009, 877, 887–891.

[29] Aloy, P., Russell, R. B., Ten thousand interactions for the molecular biologist. Nat Biotechnol 2004, 22, 1317–1321.

[30] Gavin, A. C., Aloy, P., Grandi, P., Krause, R., et al., Proteome survey reveals modularity of the yeast cell machinery. Nature 2006, 440, 631–636.

[31] Andersson, K., Arner, P., Systemic nicotine stimulates human adipose tissue lipolysis through local cholinergic and catecholaminergic receptors. Int J Obes Relat Metab Disord 2001, 25, 1225–1232.

[32] Bajaj, M., Nicotine and insulin resistance: when the smoke clears. Diabetes 2012, 61, 3078–3080.

[33] Zhang, J., Hashmi, S., Cheema, F., Al-Nasser, N., et al., Regulation of lipoprotein assembly, secretion and fatty acid beta-oxidation by Kruppel-like transcription factor, klf-3. J Mol Biol 2013, 425, 2641–2655.

[34] Bakke, J., Haj, F. G., Protein-tyrosine phosphatase 1B substrates and metabolic regulation. Semin Cell Dev Biol 2014.

[35] Murphy, C. T., Hu, P. J., Insulin/insulin-like growth factor signaling in *C. elegans*. WormBook 2013, 1–43.

[36] Padmanabhan, S., Mukhopadhyay, A., Narasimhan, S. D., Tesz, G., et al., A PP2A regulatory subunit regulates *C. elegans* insulin/IGF-1 signaling by modulating AKT-1 phosphorylation. Cell 2009, 136, 939–951.

[37] West, K. A., Brognard, J., Clark, A. S., Linnoila, I. R., et al., Rapid Akt activation by nicotine and a tobacco carcinogen modulates the phenotype of normal human airway epithelial cells. J Clin Invest 2003, 111, 81–90.

[38] Schulz, T. J., Zarse, K., Voigt, A., Urban, N., et al., Glucose restriction extends *Caenorhabditis elegans* life span by inducing mitochondrial respiration and increasing oxidative stress. Cell Metab 2007, 6, 280–293.

[39] Cypser, J. R., Johnson, T. E., Multiple stressors in *Caenorhabditis elegans* induce stress hormesis and extended longevity. J Gerontol A Biol Sci Med Sci 2002, 57, B109–114.

[40] Sobkowiak, R., Musidlak, J., Lesicki, A., In vitro genoprotective and genotoxic effect of nicotine on human leukocytes evaluated by the comet assay. Drug Chem Toxicol 2014, 37, 322–328.

[41] Calabrese, V., Mancuso, C., Ravagna, A., Perluigi, M., et al., In vivo induction of heat shock proteins in the substantia nigra following L-DOPA administration is associated with increased activity of mitochondrial complex I and nitrosative stress in rats: regulation by glutathione redox state. J Neurochem 2007, 101, 709–717.

[42] Pogocki, D., Ruman, T., Danilczuk, M., Celuch, M., Walajtys-Rode, E., Application of nicotine enantiomers, derivatives and analogues in therapy of neurodegenerative disorders. Eur J Pharmacol 2007, 563, 18–39.

[43] Hong-Qi, Y., Zhi-Kun, S., Sheng-Di, C., Current advances in the treatment of Alzheimer’s disease: focused on considerations targeting Abeta and tau. Transl Neurodegener 2012, 1, 21.

[44] Li, X., Buxbaum, J. N., Transthyretin and the brain re-visited: is neuronal synthesis of transthyretin protective in Alzheimer’s disease? Mol Neurodegener 2011, 6, 79.

[45] Kellett, K. A., Hooper, N. M., Prion protein and Alzheimer’s disease. Prion 2009, 3, 190–194.

[46] Link, C. D., Expression of human beta-amyloid peptide in transgenic *Caenorhabditis elegans*. Proc Natl Acad Sci U S A 1995, 92, 9368–9372.

[47] Shoop, R. D., Yamada, N., Berg, D. K., Cytoskeletal links of neuronal acetylcholine receptors containing alpha 7 subunits. J Neurosci 2000, 20, 4021–4029.

[48] Antolik, C., Catino, D. H., O’Neill, A. M., Resneck, W. G., et al., The actin binding domain of ACF7 binds directly to the tetratricopeptide repeat domains of rapsyn. Neuroscience 2007, 145, 56–65.

[49] Almedom, R. B., Liewald, J. F., Hernando, G., Schultheis, C., et al., An ER-resident membrane protein complex regulates nicotinic acetylcholine receptor subunit composition at the synapse. EMBO J 2009, 28, 2636–2649.

[50] Hake, S. B., Allis, C. D., Histone H3 variants and their potential role in indexing mammalian genomes: the “H3 barcode hypothesis”. Proc Natl Acad Sci U S A 2006, 103, 6428–6435.

[51] Renthal, W., Nestler, E. J., Chromatin regulation in drug addiction and depression. Dialogues Clin Neurosci 2009, 11, 257–268.

[52] Nestler, E. J., Epigenetic mechanisms of drug addiction. Neuropharmacology 2014, 76 Pt B, 259–268.

[53] Towbin, B. D., Gonzalez-Aguilera, C., Sack, R., Gaidatzis, D., et al., Step-wise methylation of histone H3K9 positions heterochromatin at the nuclear periphery. Cell 2012, 150, 934–947.

[54] Binzak, B. A., Wevers, R. A., Moolenaar, S. H., Lee, Y. M., et al., Cloning of dimethylglycine dehydrogenase and a new human inborn error of metabolism, dimethylglycine dehydrogenase deficiency. Am J Hum Genet 2001, 68, 839–847.

